# METTL1/WDR4-mediated m^7^G Hypermethylation of SCLT1 mRNA Promotes Gefitinib Resistance in Non-small Cell Lung Cancer

**DOI:** 10.1101/2025.03.26.645570

**Authors:** Shaoxuan Zhou, Yueqin Wang, Jingyao Wei, Ke An, Yong Shi, Xuran Zhang, Han Wang, Luyao Feng, Lulu He, Yu Zhang, Tong Ren, Ouwen Li, Yun-Gui Yang, Quancheng Kan, Xin Tian

## Abstract

Epidermal growth factor receptor tyrosine kinase inhibitors (EGFR-TKIs) have produced durable complete responses, but the eventual development of acquired resistance presents a huge challenge in the treatment of non-small cell lung cancer (NSCLC). N7-methylguanosine (m^7^G), a prevalent post-transcriptional modification within RNA, play regulatory roles in RNA stability, expression dynamics, and functional diversity. Despite these insights, the contribution of m^7^G methylation to EGFR-TKIs resistance remains poorly characterized. Here, we identified substantial upregulation of mRNA internal m^7^G modifications and their associated methyltransferase complex composed of methyltransferase-like 1 (METTL1) /WD repeat domain 4 (WDR4) were significantly elevated in NSCLC specimens, correlating with therapeutic resistance. Functional assays confirmed that METTL1/WDR4 enhances gefitinib resistance in both cellular and animal models via the internal RNA m^7^G methyltransferase activity in NSCLC. Mechanistically, m^7^G MeRIP-seq coupled with RNA-seq identified sodium channel and clathrin linker 1 (SCLT1) as the m^7^G target of METTL1/WDR4. METTL1/WDR4 knockdown may result in a decrease in both the demethylation level and mRNA stability of the SCLT1 transcript, while overexpression of wild-type METTL1, but not its catalytically inactive mutant, rescues mRNA stability. METTL1/WDR4-mediated m^7^G modification of SCLT1 regulates gefitinib resistance by activating the NF-κB signaling. These results establish aberrant mRNA internal m^7^G modification as a novel resistance mechanism and propose therapeutic targeting of the METTL1/WDR4-SCLT1-NF-κB axis cascade to overcome EGFR-TKIs resistance.

## Introduction

Lung cancer represents one of the most widespread malignant neoplasms globally, posing a significant threat to human health [1]. More than 85% of pulmonary malignancies fall under the non-small cell lung cancer (NSCLC) [2]. Clinical statistics indicate that nearly 60% of individuals receive diagnoses at advanced or metastatic disease, accompanied by a projected 5-year survival probability of merely 15.9% [3]. Epidermal growth factor receptor (EGFR) mutation is one of the common driving factors in the development of lung cancer [4, 5]. EGFR-tyrosine kinase inhibitors (EGFR-TKIs) constitute the primary therapeutic intervention for NSCLC patients harboring EGFR mutations [6, 7]. When compared with traditional chemotherapy regimens, EGFR-TKIs demonstrate superior efficacy in extending progression-free intervals and overall survival durations while exhibiting reduced incidence of severe adverse reactions [8]. Nevertheless, the prolonged use of EGFR-TKIs in clinical practice has its drawbacks, leading to diminished effectiveness and the eventual development of drug resistance [9, 10]. Reportedly, EGFR-TKI resistance primarily involves genetic alterations, including gene mutations, amplifications, and activation of bypass pathways [11–13]. These mechanisms are insufficient to explain the clinical emergence of drug resistance. Therefore, exploring new mechanisms and identifying new targets to reverse drug resistance will promote the development of clinical diagnostics and treatment for NSCLC.

N7-methylguanosine (m^7^G) modification, commonly found at the 5’ cap of mRNAs and within tRNAs and rRNAs, also occurs internally within mRNAs [14]. The m^7^G-cap is well-documented for its crucial roles in pre-mRNA processing, including splicing, stability, and efficient protein synthesis [15, 16]. Additionally, m^7^G at position 46 of tRNAs is a highly conserved modification crucial for regulating tRNA stability and cellular proliferation [17]. m^7^G modification is mediated by the catalytic components of the methyltransferase complex, methyltransferase-like 1 (METTL1) and WD repeat domain 4 (WDR4) [18]. Functioning as the enzymatic core, METTL1 facilitates the transfer of methyl groups derived from S-adenosylmethionine (SAM) to the N7 site of specific guanosine (G) residues within RNA substrates, thereby generating m^7^G modifications. WDR4, as the cofactor, binds to METTL1 to stabilize the complex and enhance substrate recognition, thereby enhancing methylation efficiency [14]. Within the variable loop of tRNA, the METTL1/WDR4 complex is chiefly responsible for catalyzing m^7^G methylation at the G46 site. This process stabilizes the structural integrity of tRNA and aids in achieving proper tertiary folding, which is crucial for its functional efficacy [19]. In mRNA, this complex functions as a writer enzyme, playing a crucial role in translation initiation and maintaining mRNA stability [20, 21]. These enzymes regulate the development of multiple cancer types, including lung cancer [22], liver cancer [23], head and neck tumors [24], colorectal cancer [25], bladder cancer [26], esophageal squamous cell carcinoma [27], and breast cancer [28]. Accumulating evidence suggests that abnormal m^7^G modifications influence tumor drug resistance. In hepatocellular carcinoma, the upregulation of METTL1/WDR4 in drug-resistant models contributes to lenvatinib [29] or sorafenib [30] resistance by regulating tRNA m^7^G modification and inducing downstream translational dysregulation. However, the functions and precise mechanism of mRNA internal m^7^G modification in regulating EGFR-TKIs resistance in NSCLC are not yet identified.

This study investigates the mechanisms of gefitinib resistance in NSCLC through mining public databases, performing cell sequencing, and establishing secondary drug resistance models. A comprehensive analysis of transcriptome and m^7^G MeRIP-seq data identified the METTL1/WDR4-SCLT1-NF-κB signaling axis mediated by m^7^G modification. Furthermore, the role of METTL1/WDR4-SCLT1-NF-κB signaling axis in gefitinib resistance in NSCLC was verified both *in vitro* and *in vivo*.

## Results

### High internal m^7^G modification and METTL1/WDR4 expression are associated with poor prognosis and gefitinib resistance in NSCLC

To elucidate the role of mRNA internal m^7^G modification in gefitinib resistance, we first examined the association between m^7^G modification dynamics and NSCLC progression. Bioinformatic analysis of The Cancer Genome Atlas (TCGA) revealed that significant upregulation of METTL1 and WDR4 transcripts in NSCLC tissues relative to normal samples (**Figure 1A** and B). Meanwhile, the expression levels of METTL1 and WDR4 were higher in all four histologic stages of NSCLC compared to the normal cohort (Figure 1C and D). Elevated levels of METTL1 and WDR4 exhibited a strong correlation with decreased patient survival rates (Figure 1E and F). Moreover, METTL1 and WDR4 expression showed a strong positive association with genes linked to EGFR-TKIs resistance, but a negative correlation with tumor suppressor genes (Figure 1G and H). Furthermore, METTL1 and WDR4 protein levels were markedly elevated in gefitinib-resistant clinical samples compared to gefitinib-sensitive tissues (Figure 1I and J). To confirm the connection between m^7^G modification and gefitinib resistance, we developed acquired gefitinib-resistant cell lines (PC-9/GR and HCC4006/GR). The results revealed that mRNA internal m^7^G methylation level was obviously higher in resistant cells (Figure 1K and L). Consistently, METTL1 and WDR4 protein expression were augmented in gefitinib-resistant cell lines (Figure 1M). Notably, gefitinib treatment significantly downregulated the mRNA and protein levels of METTL1 and WDR4 in sensitive cells; however, these levels remained stable in resistant cells (Figure 1N and O). These results indicated that the increased m^7^G modification and METTL1/WDR4 in gefitinib-resistant tissues and cell lines could be involved in mediating acquired gefitinib resistance in NSCLC.

**Figure 1.**
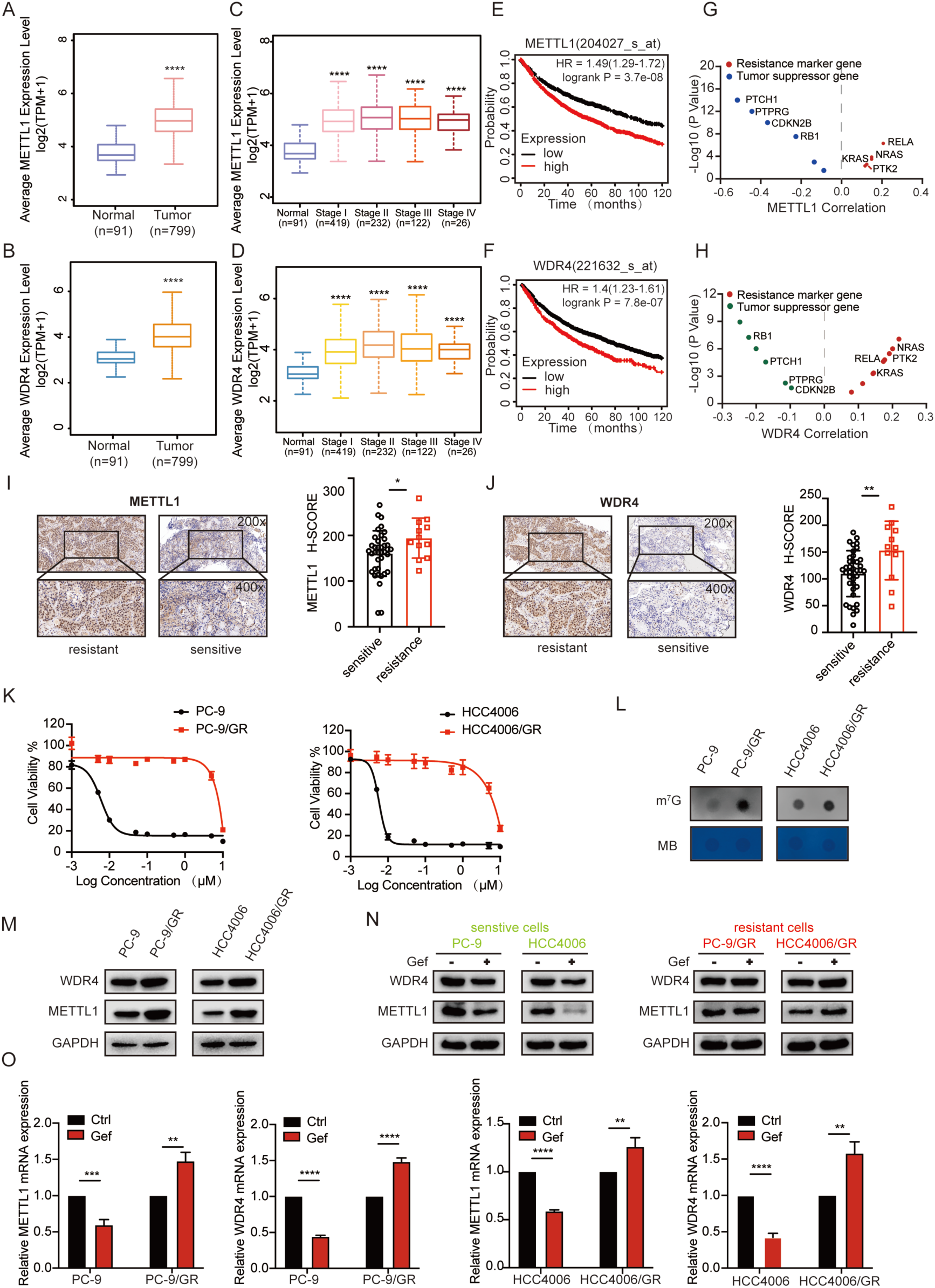
m^7^G hypermethylation and the stable expression of METL1/WDR4 are correlated to gefitinib resistance in NSCLC. **A.-B.** The expression of METTL1/WDR4 in normal tissues and non-small cell lung cancer (NSCLC) tissues from The Cancer Genome Atlas (TCGA) dataset. **C.-D.** The expression of METTL1/WDR4 in normal tissues and different individuals of NSCLC tissues from the TCGA dataset. **E.-F.** The association between the expression levels of METTL1/WDR4 and the overall survival of patients with lung cancer was assessed using the Kaplan-Meier Plotter database (http://kmplot.com/analysis/). **G.-H.** Analysis of gene expression correlation between METTL1/WDR4 and drug resistance markers or tumor suppressors was conducted using the TCGA-NSCLC dataset. **I.-J.** Representative images of METTL1/WDR4 IHC staining and the quantitative H-scores gefitinib-sensitive (n = 38) or acquired gefitinib-resistant NSCLC patients (n = 12). Image magnification: 200x (upper panel) and 400x (lower panel). **K.** Sensitive (PC-9 and HCC4006) and acquired gefitinib-resistant (PC-9/GR and HCC4006/GR) cells were subjected to treatment with varying concentrations of gefitinib for 72 hours, and the half-maximal inhibitory concentration (IC_50_) was determined using the Cell Counting Kit-8 (CCK-8) assay. **L.** The m^7^G level in sensitive and acquired gefitinib-resistant cells was detected by dot blot assay. **M.** The expression level of METTL1/WDR4 in sensitive and acquired gefitinib-resistant cells was detected by Western blot assay. **N.-O.** Sensitive and resistant cells were exposed to gefitinib (1 µM) for 24 hours. The protein expression levels of METTL1/WDR4 were evaluated using Western blotting (**N**), while mRNA expression levels were assessed by qRT-PCR (**O**). Data are shown as mean ± SD. *, *P* < 0.05; **, *P* < 0.01; ***, *P* < 0.001; ****, *P* < 0.0001. An unpaired t-test was used unless otherwise stated.

### METTL1/WDR4 knockdown modulates cell proliferation, apoptosis, and gefitinib resistance

To investigate the functional impact of METTL1/WDR4-regulated m^7^G methylation on gefitinib resistance mechanisms in NSCLC, we silenced the expression of METTL1/WDR4 using small interfering RNA (siRNA) (**Figure 2A** and B, Figure S1A and B). METTL1/WDR4 knockdown inhibited cell proliferation of PC-9/GR and HCC4006/GR cells (Figure 2C, Figure S1C). Flow cytometry analysis demonstrated that METTL1/WDR4 knockdown increased cell apoptosis and this effect was further enhanced by gefitinib (Figure 2D, Figure S1D). Western blot analysis detected that knocking down of METTL1/WDR4 enhanced caspase-3 cleavage in resistant cells (Figure 2E, Figure S2E). Next, we generated stable METTL1 and WDR4 knockdown cells using short hairpin RNA (shRNA). Knockdown of METTL1 or WDR4 expression markedly reduced cell proliferation and colony formation, triggered apoptosis and enhanced gefitinib sensitivity in resistant cells. The observed phenotypic changes could be rescued by stable overexpression of wild-type METTL1 or WDR4, whereas the catalytically deficient METTL1 mutant failed to restore these effects (Figure 2F–K, Figure S1F and G). These findings indicate that METTL1/WDR4 critically contributes to the development of acquired gefitinib resistance.

**Figure 2.**
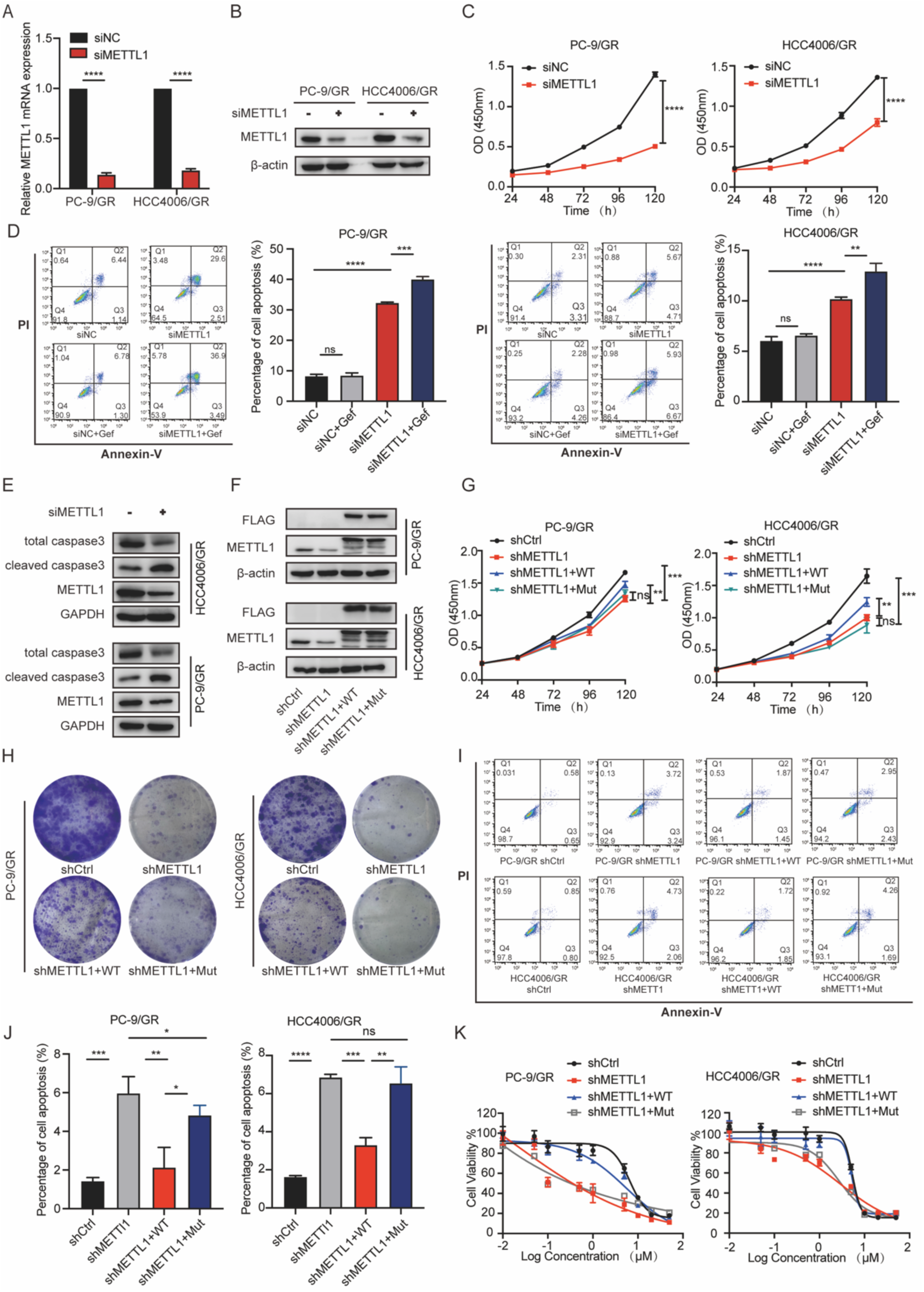
Inhibition of METTL1 overcomes gefitinib resistance *in vitro*. **A.-B.** The mRNA (**A**) and protein expression (**B**) levels of METTL1 in PC-9/GR and HCC4006/GR cells, treated with siRNA for 72 hours, were evaluated using qRT-PCR and Western blotting, respectively. **C.** Knockdown of METTL1 using siRNA was performed in PC-9/GR and HCC4006/GR cells, and the cell viability was assessed using the CCK-8 assay. **D.** PC-9/GR and HCC4006/GR cells transfected with either siNC or siMETTL1 were exposed to gefitinib (1 µM) for 72 hours, and cell apoptosis was assessed using flow cytometry. Bar graphs were generated to quantify the data obtained from Annexin V-FITC/PI staining. **E.** The apoptosis marker Cleaved Caspase 3 was detected by Western blotting in PC-9/GR and HCC4006/GR cells treated with siNC or siMETTL1 for 72 hours. **F.** The protein expression of METTL1 was assessed in PC-9/GR and HCC4006/GR cells transfected with shMETTL1, wild-type METTL1 (METTL1-WT), or the catalytically inactive mutant (METTL1-Mut, EIR/AAA) using Western blotting analysis. **G.-H.** The Cell proliferation was detected by CCK-8 (**G**) or colony formation (**H**) assay in PC-9/GR and HCC4006/GR cells transfected with shMETTL1, METTL1-WT, or METTL1-Mut. **I.-J.** PC-9/GR and HCC4006/GR cells transfected with shMETTL1, METTL1-WT, or METTL1-Mut were analyzed for cell apoptosis (**I**) using flow cytometry. Bar graphs (**J**) were generated to quantify the data obtained from Annexin V-FITC/PI staining. **K.** PC-9/GR and HCC4006/GR cells transfected with shMETTL1, METTL1-WT, or METTL1-Mut were treated with varying concentrations of gefitinib for 72 hours. The cell viability was assessed using the CCK-8 assay. Data are shown as mean ± SD. ns, not significant; *, *P* < 0.05; **, *P* < 0.01; ***, *P* < 0.001; ****, *P* < 0.0001. An unpaired t-test was used unless otherwise stated.

### METTL1 knockdown inhibits tumor growth in acquired gefitinib-resistant cell-derived xenograft model

To further assess the impact of m^7^G modification to gefitinib resistance *in vivo*. Tumor xenografts were established by subcutaneously injecting PC-9/GR cells stably expressing control shRNA, METTL1 shRNA or METTL1 shRNA rescued with WT METTL1 into the flank of BALB/c nude mice. Consistent with our *in vitro* results, METTL1 knockdown significantly reduced both tumor size and tumor weight when compared to the control group. This effect was reversed by stable overexpression of WT METTL1 (**Figure 3A–C**). The tumors obtained from the mice bearing shMETTL1 cells showed less proliferation as indicated by a significant reduction of Ki67 staining. Wild-type METTL1 overexpression efficiently rescued the cell proliferation inhibition induced by the knockdown of METTL1 (Figure 3D and E). These data suggested that METTL1 plays a critical role in acquired gefitinib resistance.

**Figure 3.**
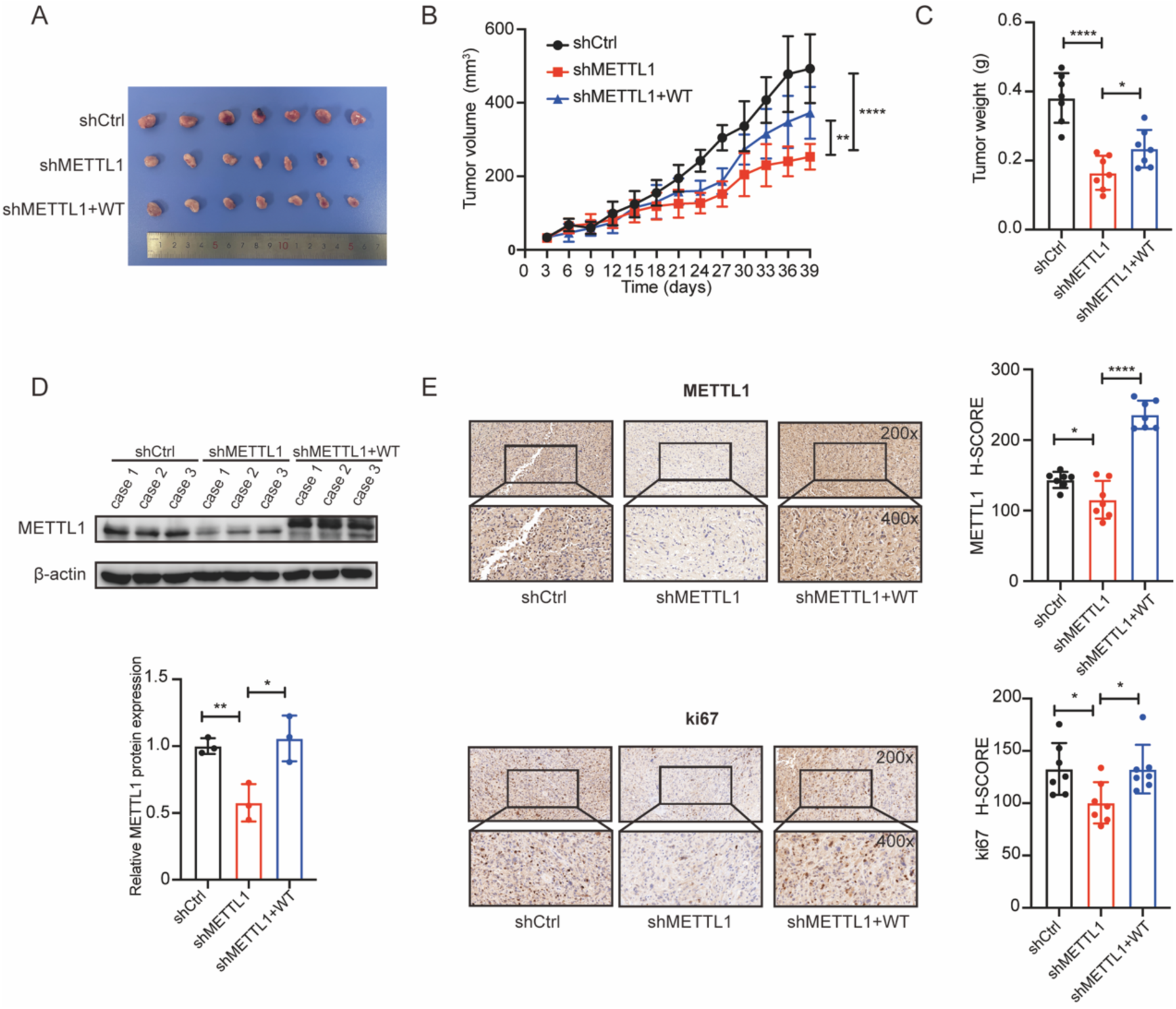
Knockdown of METTL1 inhibits tumor growth *in vivo*. **A.** Overview of tumors in BALB/c nude mice subcutaneously implanted with shCtrl, shMETTL1, and rescue METTL1 PC-9/GR cells (n = 7). **B.-C.** Mean volumes (**B**) and tumor weights (**C**) of tumor xenografts obtained from BALB/c nude mice subcutaneously implanted with shCtrl, shMETTL1, and rescue METTL1 PC-9/GR cells (n = 7). **D.** Western blot analysis of METTL1 expression in tumor tissues from the shCtrl, shMETTL1, and rescue METTL1 groups (n = 3). **E.** Representative images displaying METTL1 and Ki67 immunohistochemical staining, alongside quantitative H-scores of tumors obtained from the shCtrl, shMETTL1, and rescue METTL1 groups xenografts, are presented. Image magnification: 200× (upper panel) and 400× (lower panel). Data are shown as mean ± SD. *, *P* < 0.05; **, *P* < 0.01; ****, *P* < 0.0001. An unpaired t-test was used unless otherwise stated.

### METTL1/WDR4 upregulation mediates m^7^G hypermethylation in gefitinib resistant cells

Given that METTL1/WDR4 was identified as the m^7^G writers, we performed the m^7^G dot blot assay and found that the knockdown of METTL1/WDR4 in PC-9/GR cells decreased the m^7^G modification level of mRNA, which could be rescued by wild-type METTL1 or WDR4, but not the catalytic mutant METTL1 (**Figure 4A**, Figure S2A and B). To explore the potential mechanism of mRNA internal m^7^G modification promoting gefitinib resistance, we further employed m^7^G-modified RNA immunoprecipitation sequencing (MeRIP-seq) following the removal of the 5’ cap structure from mRNA, alongside RNA sequencing (RNA-seq) analysis (Figure 4B). MeRIP-seq revealed that the majority of m^7^G peaks were concentrated within the coding sequence (CDS) region (Figure 4C). Hypergeometric Optimization of Motif EnRichment (HOMER) analyzed that the GANGAN modification motif was highly enriched within m^7^G sites in PC-9/GR with METTL1/WDR4 knockdown and the negative control (Figure 4D). MeRIP-seq revealed that METTL1 knockdown resulted in a decreased abundance of m^7^G peaks in 814 transcripts and increased abundance in 84 transcripts, while WDR4 knockdown induced upregulation of m^7^G peaks in 76 transcripts and downregulation of m^7^G peaks in 704 transcripts (Figure 4E). To uncover gefitinib resistance-related genes controlled by METTL1/WDR4, gene expression profiled by RNA-seq and transcripts having m^7^G sites identified by MeRIP-seq were co-analyzed. There were 27 genes that showed m^7^G hypomethylation in shMETTL1/WDR4 cells and exhibited not only decreased expression in sensitive cells (PC-9 + Gef) but also stabilized expression in resistant cells (PC-9/GR + Gef) upon gefitinib treatment (Figure 4F; Table S1 and S2). Next, we chose 11 genes as potential targets of METTL1/WDR4-mediated m^7^G alteration because they were demonstrated to be linked to drug resistance. Further studies with qRT-PCR demonstrated that METTL1/WDR4 knockdown markedly reduced mRNA expression of SCLT1 in both PC-9/GR and HCC4006/GR cells (Figure 4G). As for the other genes of interest, we also performed validation (Figure S3A–D). Collectively, these findings indicated that SCLT1 might be the key gene targeted by METTL1/WDR4-mediated m^7^G modification involved in gefitinib resistance.

**Figure 4.**
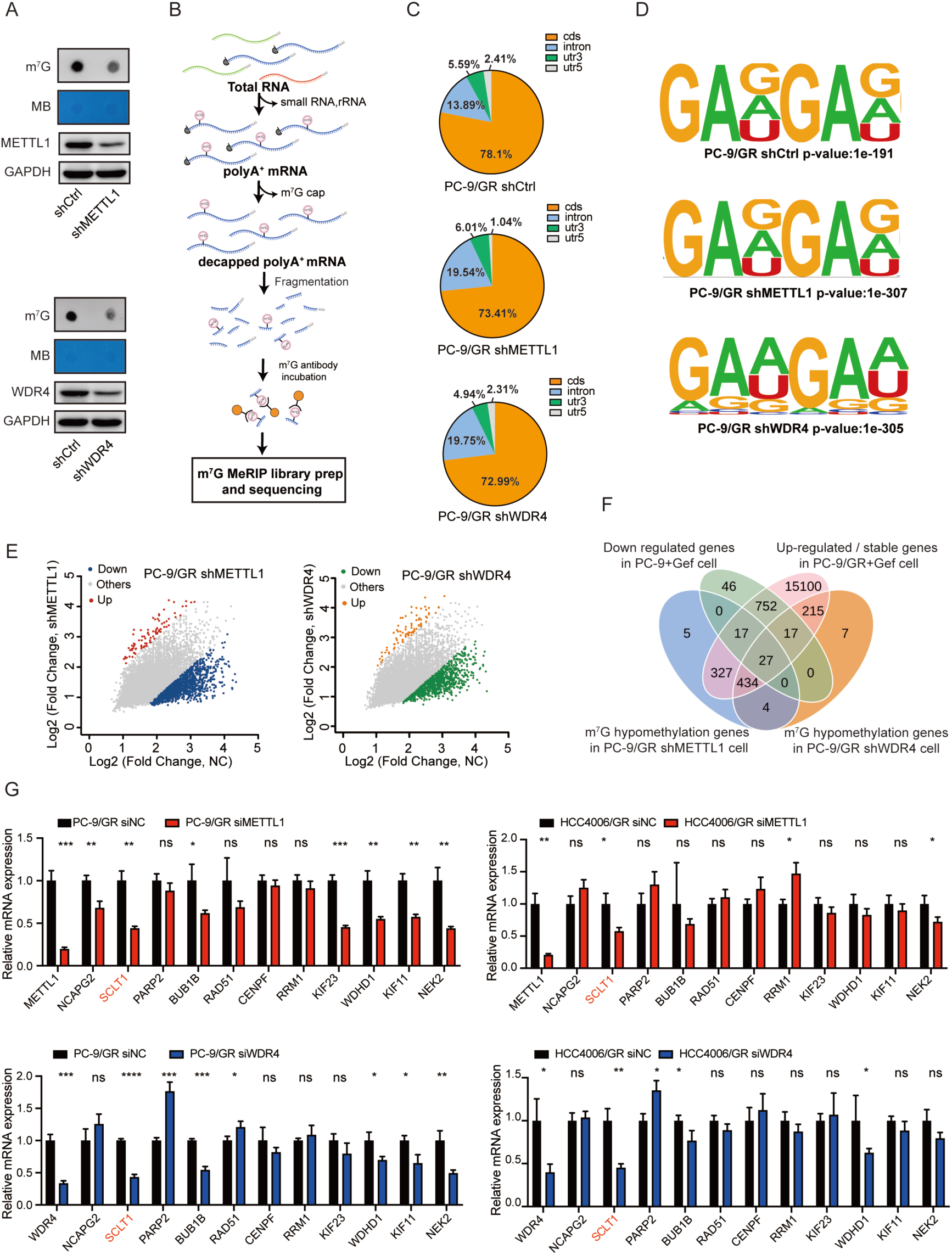
Knockdown of METTL1/WDR4 promotes m^7^G demethylation in gefitinib-resistant NSCLC cells. **A.** The m^7^G level in PC-9/GR cells after the knockdown of METTL1/WDR4 was detected by dot blot assay. **B.** The flowchart for m^7^G MeRIP-seq. **C.** The pie chart displays the percentage of m^7^G sites within RNA categories and their distribution on mRNA. **D.** Sequence motif was identified from MeRIP-seq analysis in PC-9/GR cells after knockdown of METTL1/WDR4. **E.** The scatter plot depicts significant changes in gene expression observed in PC-9/GR shMETTL1 and PC-9/GR shWDR4 cells. **F.** Venn diagram depicting potential candidates for m^7^G modification among METTL1/WDR4 target genes. **G.** qRT-PCR was conducted to assess the mRNA levels of target genes in PC-9/GR and HCC4006/GR cells after the knockdown of METTL1/WDR4. Data are shown as mean ± SD. ns, not significant; *, *P* < 0.05; **, *P* < 0.01; ***, *P* < 0.001; ****, *P* < 0.0001. An unpaired t-test was used unless otherwise stated.

### METTL1/WDR4-mediated m^7^G modification regulates SCLT1 mRNA stability

To further determine the mechanism underlying METTL1/WDR4-mediated m^7^G modification in the regulation of SCLT1, comprehensive analyses were conducted. Integrative Genomics Viewer (IGV) analysis showed that METTL1/WDR4 knockdown reduced m^7^G deposition within the CDS region of SCLT1, accompanied by diminished transcriptional abundance of *SCLT1* in PC-9/GR cells (**Figure 5A**). MeRIP-qPCR validation experiments further corroborated that METTL1/WDR4 knockdown decreased the m^7^G modification abundance of the CDS region of SCLT1 mRNA in PC-9/GR and HCC4006/GR cells (Figure 5B). We then investigated whether METTL1/WDR4 knockdown affected the stability of SCLT1 mRNA. As shown in Figure 5C, after actinomycin D treatment, the half-life of SCLT1 mRNA was notably decreased upon METTL1/WDR4 suppression in PC-9/GR and HCC4006/GR cells. Rescue experiments demonstrated that reintroduction of wild-type METTL1, rather than the catalytically inactive mutant, restored both SCLT1 protein expression and mRNA stability (Figure 5D and E). These results supported that METTL1/WDR4-mediated m^7^G hypermethylation regulates SCLT1 expression by maintenance of SCLT1 mRNA stability. Subsequent investigations evaluated the clinical significance of SCLT1 in gefitinib resistance in NSCLC. Bioinformatic interrogation of TCGA database demonstrated positive associations between SCLT1 expression and EGFR-TKIs resistance markers, contrasted by inverse correlation with tumor suppressor genes (Figure 5F). Moreover, immunohistochemical analysis identified that the SCLT1 protein level was significantly higher in gefitinib-resistant clinical samples than in gefitinib-sensitive tissues (Figure 5G). In addition, high expression of SCLT1 was significantly associated with poor patient survival (Figure 5H). Taken together, our findings demonstrated that SCLT1 functioned as the key downstream target of METTL1/WDR4-mediated m^7^G modification in acquired gefitinib resistance in NSCLC.

**Figure 5.**
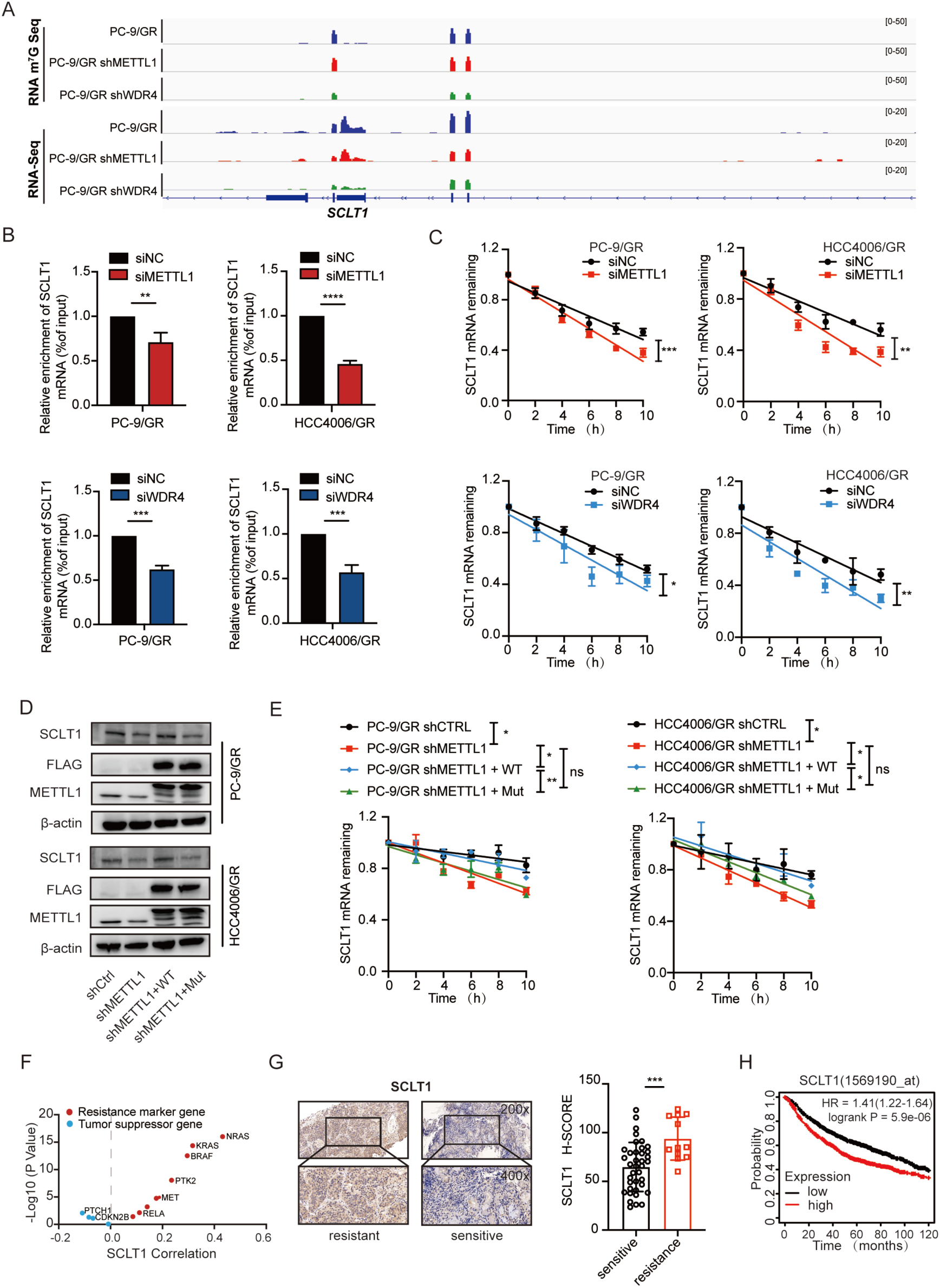
METTL1/WDR4-mediated m^7^G hypermethylation of SCLT1 mRNA maintains its stability. **A.** IGV analysis revealed alterations in mRNA expression and m^7^G levels of SCLT1 in PC-9/GR cells following the knockdown of METTL1 and WDR4. **B.** Decapped mRNA, purified and subjected to immunoprecipitation using an anti-m^7^G antibody, underwent qRT-PCR analysis to assess m^7^G levels of SCLT1 in PC-9/GR and HCC4006/GR cells. **C.** After 72 hours of treatment with siMETTL1 or siWDR4, PC-9/GR and HCC4006/GR cells were exposed to actinomycin-D (5 μg/mL) for various time intervals. Subsequently, qRT-PCR was employed to analyze the mRNA half-life of SCLT1. **D.** The protein expression of METTL1 and SCLT1 was assessed in PC-9/GR and HCC4006/GR cells transfected with shMETTL1, METTL1-WT, or METTL1-Mut using Western blotting analysis. **E.** qRT-PCR was employed to analyze the mRNA half-life of SCLT1 in PC-9/GR and HCC4006/GR cells transfected with shMETTL1, METTL1-WT, or METTL1-Mut. **F.** Analysis of gene expression correlation between SCLT1 and drug resistance markers or tumor suppressors was conducted using the TCGA-NSCLC dataset. **G.** Representative images of SCLT1 staining and the quantitative H-scores gefitinib-sensitive (n = 38) or gefitinib-resistant NSCLC patients (n = 12). Image magnification: 200x (upper panel) and 400x (lower panel). **H.** The association between the expression levels of SCLT1 and the overall survival of patients with lung cancer was assessed using the Kaplan-Meier Plotter database. Data are shown as mean ± SD. ns, not significant; *, *P* < 0.05; **, *P* < 0.01; ***, *P* < 0.001; ****, *P* < 0.0001. An unpaired t-test was used unless otherwise stated.

### SCLT1 knockdown hinders the proliferation and gefitinib-resistance *in vitro* and *in vivo*

To investigate the involvement of SCLT1 in gefitinib resistance, SCLT1 expression was found to be up-regulated in both gefitinib-resistant PC-9/GR and HCC4006/GR cells (**Figure 6A**). Meanwhile, SCLT1 was also found to be up-regulated in gefitinib-resistant NSCLC samples (Figure 5E). In addition, gefitinib treatment significantly downregulated the mRNA and protein levels of SCLT1 in sensitive cells; however, the levels remained stable in resistant cells (Figure 6B and C). These findings suggested a correlation between SCLT1 and gefitinib resistance in NSCLC. Functional assessments utilizing CCK-8 and clonogenic assays showed that knocking down SCLT1 significantly inhibited cell proliferation and colony formation of gefitinib-resistant cells (Figure 6D–G). Moreover, increased therapeutic responsiveness to gefitinib was evident in the SCLT1 knockdown group (Figure 6H). Flow cytometry analysis demonstrated that SCLT1 knockdown induced cell apoptosis and this effect was further enhanced by gefitinib (Figure 6I). *In vivo*, the knockdown of SCLT1 significantly reduced tumor size and tumor weight of PC-9/GR xenografts compared to the control group (Figure 6J–L). These findings indicated that SCLT1 is vital to cell proliferation, apoptosis, and gefitinib resistance in NSCLC.

**Figure 6.**
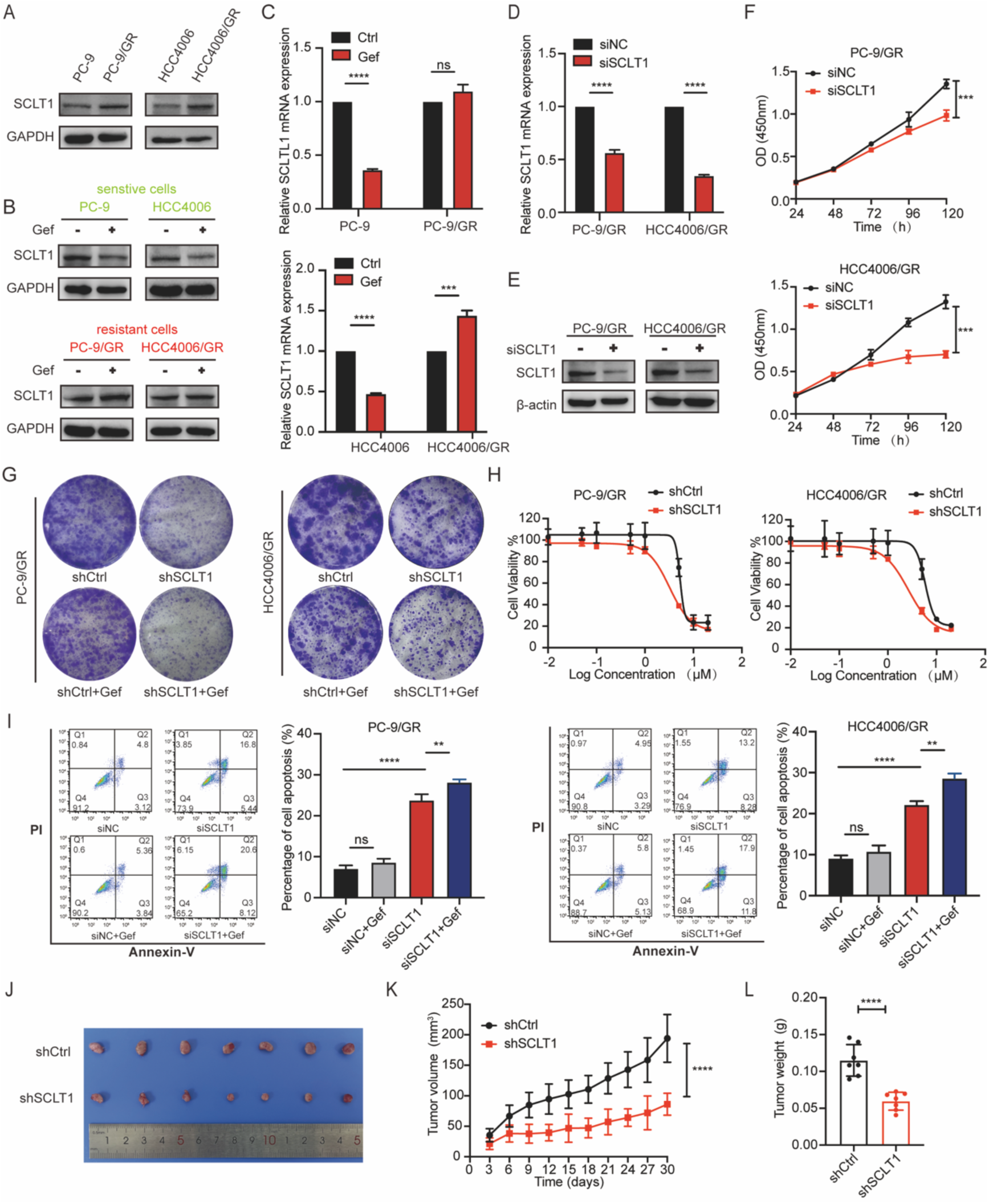
Inhibiting SCLT1 effectively overcomes gefitinib resistance both *in vitro* and *in vivo*. **A.** The expression level of SCLT1 in sensitive and acquired gefitinib-resistant cells was detected by Western blot assay. **B.-C.** Sensitive and resistant cells were exposed to gefitinib (1 µM) for 24 hours. The protein expression levels of SCLT1 were evaluated using Western blotting (**B**), while mRNA expression levels were assessed by qRT-PCR (**C**). **D.-E.** The mRNA (**D**) and protein expression (**E**) levels of SCLT1 in PC-9/GR and HCC4006/GR cells, treated with siRNA for 72 hours, were evaluated using qRT-PCR and Western blotting, respectively. **F.** Knockdown of SCLT1 using siRNA was performed in PC-9/GR and HCC4006/GR cells, and the cell viability was assessed using the CCK-8 assay. **G.** The colony formation ability was evaluated in PC-9/GR and HCC4006/GR cells transfected with shSCLT1 using the colony formation assay. **H.** PC-9/GR and HCC4006/GR cells, following transfected with shSCLT1, were exposed to varying concentrations of gefitinib for a duration of 72 hours. Subsequently, the CCK-8 assay was employed to determine cell viability. **I.** PC-9/GR and HCC4006/GR cells transfected with either siSCLT1 or siCtrl were exposed to gefitinib (1 µM) for 72 hours, and cell apoptosis was assessed using flow cytometry. Bar graphs were generated to quantify the data obtained from Annexin V-FITC/PI staining. **J.** Overview of tumors in BALB/c nude mice subcutaneously implanted with shCtrl and shSCLT1 PC-9/GR cells (n = 7). **K.-L.** Mean volumes (**K**) and tumor weights (**L**) of tumor xenografts obtained from BALB/c nude mice subcutaneously implanted with shCtrl and shSCLT1 PC-9/GR cells (n = 7). Data are shown as mean ± SD. ns, not significant; **, *P* < 0.01; ***, *P* < 0.001; ****, *P* < 0.0001. An unpaired t-test was used unless otherwise stated.

### SCLT1 overexpression partially rescues cellular phenotypes in gefitinib-resistant cells with METTL1/WDR4 knockdown

The results mentioned above showed that METTL1/WDR4 regulates gefitinib resistance by modulating the mRNA stability of SCLT1. To validate these findings, we overexpressed the downstream target SCLT1 in Gefitinib-resistant cells with METTL1/WDR4 knockdown (**Figure 7A**). Our findings demonstrate that SCLT1 overexpression partially rescues cell growth and apoptosis in METTL1/WDR4-knockdown gefitinib-resistant cells (Figure 7B–D). Moreover, SCLT1 overexpression also restores gefitinib sensitivity in these cells (Figure 7E). Together, these findings highlight the essential function of the METTL1/WDR4-SCLT1 axis in regulating gefitinib resistance.

**Figure 7.**
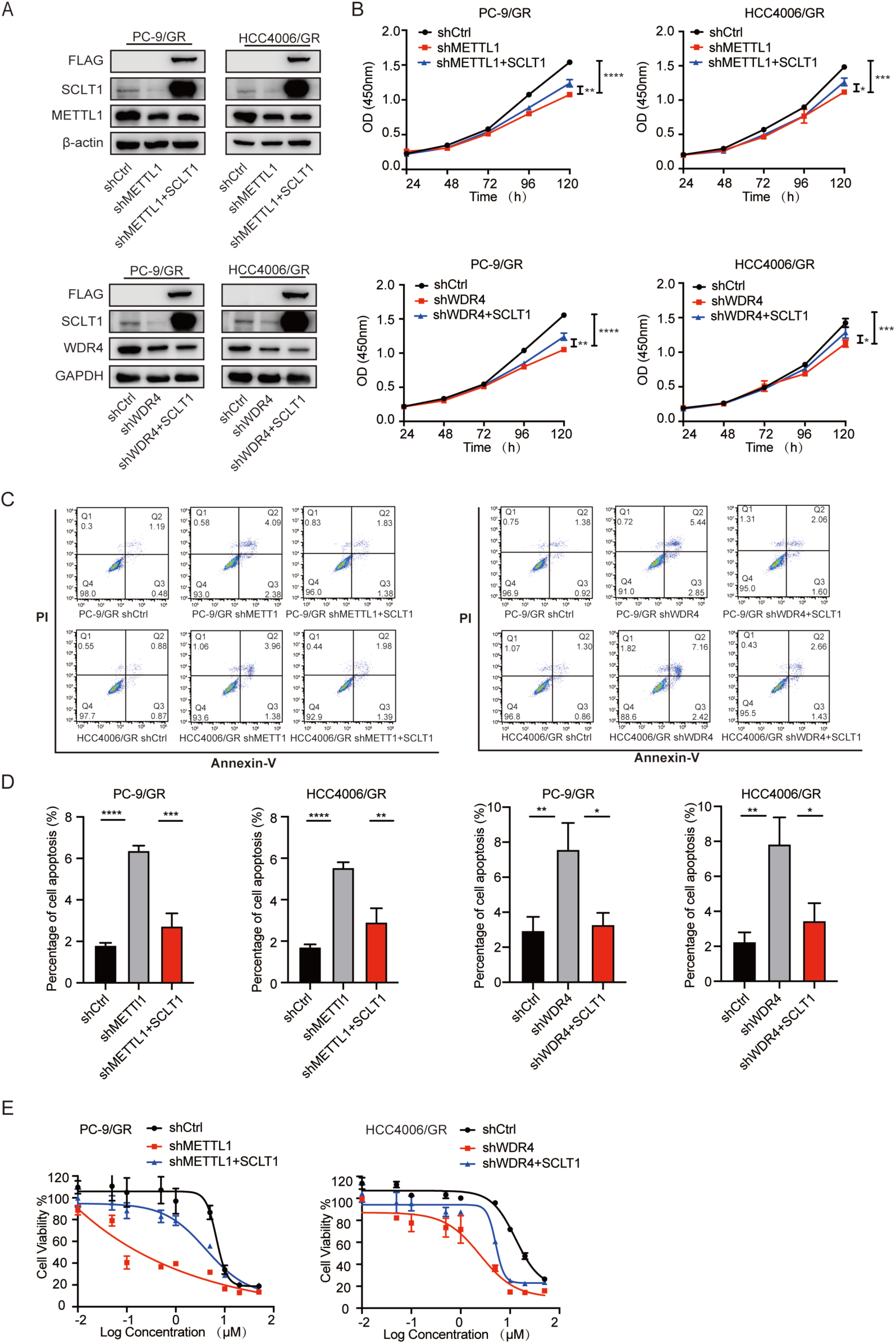
Functional restoration via SCLT1 in gefitinib-resistant cells with METTL1/WDR4 knockdown. **A.** The expression level of METTL1/WDR4 and SCLT1 in PC-9/GR and HCC4006/GR cells. **B.-D.** PC-9/GR and HCC4006/GR cells pre-treated with shMETTL1/WDR4 were stably transfected with wild type SCLT1 and cell proliferation (**B**) or cell apoptosis (**C**) was respectively detected by CCK-8 assay or Annexin V-FITC/PI staining. Bar graphs (**D**) were generated to quantify the data obtained from Annexin V-FITC/PI staining **E.** PC-9/GR and HCC4006/GR cells transfected with shMETTL1/WDR4 or rescue SCLT1 were treated with varying concentrations of gefitinib for 72 hours. The cell viability was assessed using the CCK-8 assay. Data are shown as mean ± SD. *, *P* < 0.05; **, *P* < 0.01; ***, *P* < 0.001; ****, *P* < 0.0001. An unpaired t-test was used unless otherwise stated.

### NF-κB pathway activation by METTL1/WDR4-SCLT1 underlies gefitinib resistance in NSCLC

Subsequently, we delved into the molecular mechanisms underlying METTL1/WDR4-mediated-SCLT1 m^7^G modification in gefitinib-acquired resistance in NSCLC. Gene Ontology (GO) pathway enrichment analysis identified the potential pathways influenced by the METTL1/WDR4-SCLT1 axis. Strikingly, METTL1, WDR4, or SCLT1 knockdown enriched the relevant pathways in the NF-κB and apoptosis pathways (Figures S5A–C). These findings were validated by Western blot analysis, indicating that silencing METTL1, WDR4, or SCLT1 inhibits the NF-κB pathway in resistant cells (**Figure 8A**). Additional research revealed that QNZ (EVP4593) effectively suppresses NF-κB pathway activity, leading to a decline in the proliferation of resistant cells while simultaneously triggering apoptosis (Figure 8B–F). The results underscore the central role of the METTL1/WDR4-SCLT1 axis in driving gefitinib resistance in NSCLC by modulating NF-κB signaling. Notably, suppressing NF-κB pathway activity can reverse the resistant phenotype, suggesting that targeting the METTL1/WDR4-SCLT1-NF-κB axis holds significant potential as a therapeutic approach to address resistance in NSCLC.

**Figure 8.**
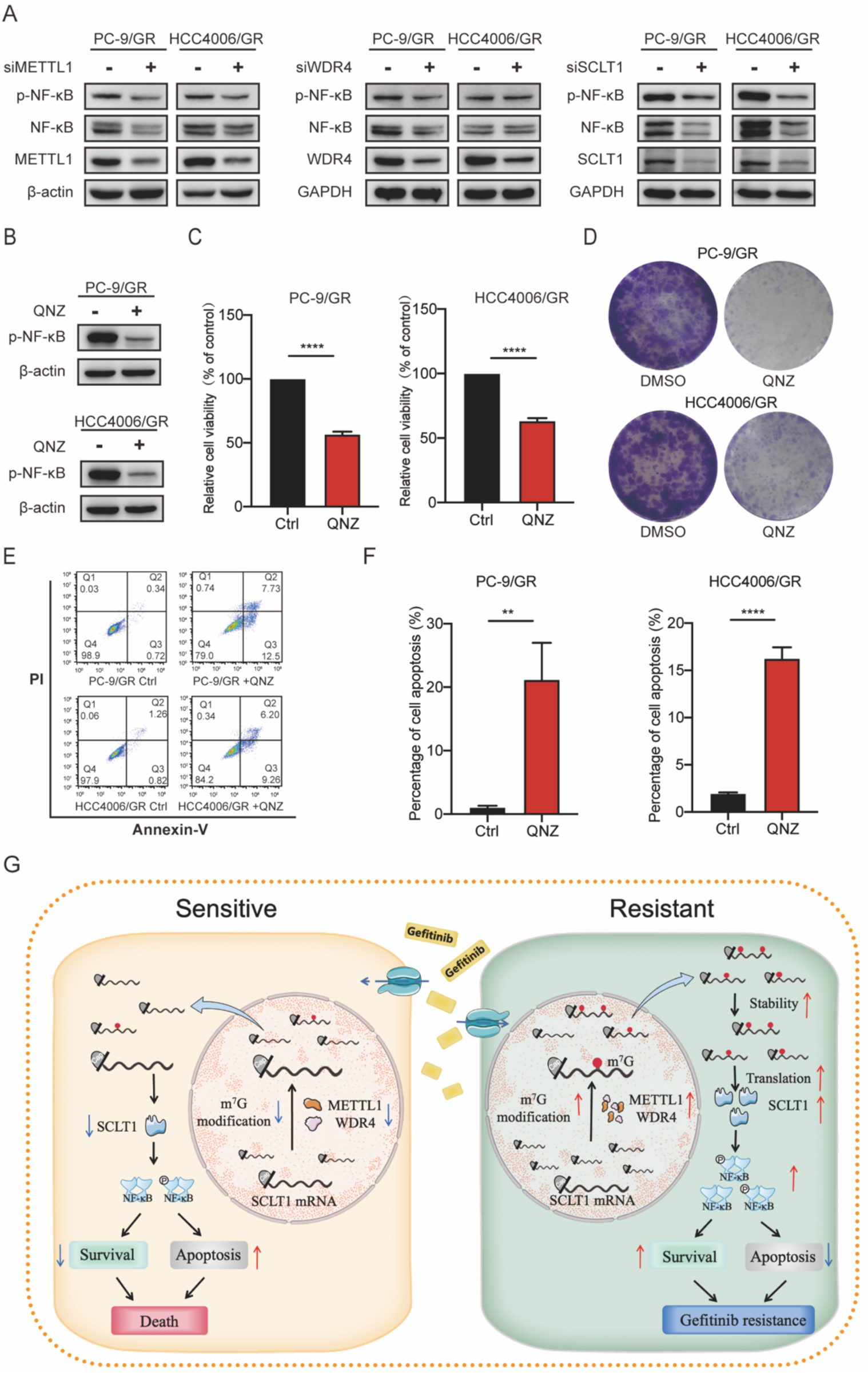
NF-κB activation mediated by METTL1/WDR4-SCLT1 confers gefitinib resistance in NSCLC. **A.** The protein expression levels of NF-κB were evaluated via Western blotting in PC-9/GR and HCC4006/GR cells following a 72-hour treatment with siMETTL1, siWDR4, or siSCLT1. **B.** The protein expression levels of p-NF-κB in PC-9/GR and HCC4006/GR cells treated with DMSO or QNZ (1 µM) for 72 hours were evaluated using Western blotting. **C.-D.** Cell proliferation was evaluated via the CCK-8 (**C**) and colony formation assays (**D**) in PC-9/GR and HCC4006/GR cells exposure to DMSO or QNZ (1 µM) for 72 hours. **E.-F.** PC-9/GR and HCC4006/GR cells were exposed to DMSO or QNZ (1 µM) for 72 hours, and cell apoptosis (**E**) was assessed using flow cytometry. Bar graphs (**F**) were generated to quantify the data obtained from Annexin V-FITC/PI staining. **G.** Proposed model illustrating the mechanism of acquired gefitinib resistance in NSCLC through aberrant m^7^G hypermethylation via the METTL1/WDR4-SCLT1-NF-κB axis. Data are shown as mean ± SD. **, *P* < 0.01; ****, *P* < 0.0001. An unpaired t-test was used unless otherwise stated.

## Discussion

Gefitinib, serving as a key targeted therapy for NSCLC patients with EGFR-activating mutations, shows remarkable clinical effectiveness in bringing about tumor regression and extending survival [31]. However, the rise of acquired resistance continues to pose a significant obstacle to achieving lasting therapeutic outcomes [32, 33]. Emerging evidence indicates that RNA epigenetic modifications, particularly m^5^C and m^6^A methylation of mRNA, play essential roles in drug resistance across multiple malignancies [34, 35]. Additionally, RNA secondary structure has been implicated in EGFR TKI resistance by controlling YRDC mRNA translation in NSCLC [36]. Our study identifies a new mechanism through which mRNA internal m^7^G modifications promote gefitinib resistance in NSCLC via the METTL1/WDR4-SCLT1-NF-κB axis in an m^7^G-dependent manner. These findings not only deepen our understanding of gefitinib resistance but also suggest that targeting m^7^G modifications could represent a promising approach to counteract resistance in NSCLC.

Our study demonstrates a profound clinical relevance of METTL1/WDR4 upregulation in gefitinib-resistant NSCLC patients (Figure 1I and J). Consistently, we observed elevated m^7^G methylation levels in resistant cells compared to sensitive counterparts (Figure 1L), suggesting a positive feedback loop between drug resistance and epigenetic modification. These findings align with preliminary evidence from liver cancer models, where lenvatinib resistance correlates with increased METTL1/WDR4 expression and RNA m^7^G methylation [29]. Similarly, the study showed that R-2HG treatment suppresses FTO expression in sensitive leukemia cells, whereas this effect is absent in resistant cells [37]. Consistent with this, we detected decreased mRNA and protein levels of METTL1 and WDR4 in gefitinib-sensitive cells, yet not in resistant cells. These results highlight the possible role of m^7^G modification in gefitinib resistance and suggest that targeting the m^7^G pathway could be a promising approach to overcome gefitinib resistance in NSCLC.

Recent investigations have mainly centered on the function of tRNA m^7^G modifications in cancer progression and drug resistance [14] with limited reports on the role of internal m^7^G modifications in mRNA within this context [38, 39]. For instance, METTL1/WDR4-mediated tRNA m^7^G modifications boost translation efficiency and fuel cancer progression in malignancies such as lung and liver cancer [22, 23]. Additionally, tRNA m^7^G modifications have been implicated in drug resistance in hepatocellular carcinoma [29, 30] and oral squamous cell carcinoma [40]. However, the role of mRNA internal m^7^G modifications in gefitinib resistance remains underexplored. Our study reveals that METTL1/WDR4-mediated internal m^7^G modifications in mRNA regulate gefitinib resistance in NSCLC by stabilizing SCLT1 mRNA. Nevertheless, the role of m^7^G modifications in other RNA types, such as tRNA, remains underexplored. This gap in knowledge may limit our complete understanding of the impact of m^7^G modifications on gefitinib resistance. Additional research is needed to elucidate the broader role of m^7^G modifications across different RNA species and their collective impact on gefitinib resistance.

SCLT1 (Sodium Channel and Clathrin Linker 1) is an adaptor protein that links SCN10A to clathrin, thereby potentially regulating SCN10A channel activity, which in turn may affect cell proliferation and migration [41]. Additionally, bioinformatics discovered a correlation between the SCLT1 gene and sensitivity to paclitaxel in lung squamous cell carcinoma [42]. In our research, we observed elevated SCLT1 levels in both gefitinib-resistant cells and NSCLC patients with gefitinib resistance. By knocking down SCLT1, we observed a reversal of gefitinib resistance both *in vitro* and *in vivo*. These results underscore the potential significance of SCLT1 in mediating resistance to gefitinib in NSCLC. However, current m^7^G MeRIP-seq technology lacks single-base resolution [8, 13], which limits our ability to generate precise point mutations in SCLT1. Despite this limitation, we conducted rescue experiments by reintroducing wild-type SCLT1, which partially restored cellular phenotypes in gefitinib-resistant cells with METTL1/WDR4 knockdown. Our study reveals that METTL1-mediated m^7^G modification of SCLT1 mRNA promotes acquired resistance to gefitinib in NSCLC.

The NF-κB signaling cascade has been previously linked to resistance against EGFR-TKI treatments [43]. In the present study, GO enrichment analysis revealed that genes downregulated upon METTL1/WDR4/SCLT1 knockdown in PC-9/GR cells were notably enriched in the NF-κB signaling. Functional experiments confirmed that METTL1/WDR4/SCLT1knockdown blocked the phosphorylation of NF-κB pathway. Furthermore, treatment with the NF-κB inhibitor QNZ notably suppressed cell proliferation and promoted apoptosis in gefitinib-resistant cells. These results demonstrate that METTL1/WDR4-SCLT1 axis regulates gefitinib resistance through NF-κB signaling.

In summary, this study uncovers a previously unrecognized mechanism involving the METTL1/WDR4-SCLT1-NF-κB axis, which plays a pivotal role in regulating acquired resistance to gefitinib in NSCLC (Figure 8G). Suppression of the METTL1/WDR4-SCLT1-NF-κB axis effectively overcomes gefitinib resistance through an m^7^G-dependent regulatory mechanism. Our research uncovers novel mechanisms of gefitinib resistance and identifies m^7^G modification as a potential therapeutic target for NSCLC.

## Materials and methods

### Patients and samples

The clinical samples utilized in this study were obtained from patients in the Department of Thoracic Surgery and Department of Oncology at the First Affiliated Hospital of Zhengzhou University. All samples underwent pathological diagnosis confirming lung cancer. According to medical records, these patients were classified as either sensitive or resistant to EGFR-TKIs drug therapy. Among all tumor samples, EGFR gene mutations consisted of exon 19 deletion (19del) or exon 21 point mutation (L858R), with no EGFR T790M mutation detected. Acquired EGFR-TKIs resistance was operationally defined as disease progression in lung cancer patients with EGFR mutations after an average duration of 10 to 14 months following treatment with EGFR-TKIs. Patients who underwent surgery, chemotherapy, radiotherapy, or immunotherapy in the six months prior were not included in this study.

### Immunohistochemistry (IHC) staining

Briefly, tissue sections were initially baked to enhance tissue adherence, followed by deparaffinization and rehydration. Antigen retrieval was performed to improve antibody binding efficiency, followed by blocking to mitigate non-specific binding. Subsequent steps included primary and secondary antibody incubations for antigen detection, DAB staining for visualization, and nuclear counterstaining for contrast enhancement. Finally, the sections underwent dehydration, air-drying, and mounting with neutral resin for preservation. Slides were scanned and analyzed using Quant Center software. The concentrations of anti-METTL1 (Catalog No. ab271063; Abcam, Cambridge, MA, US), anti-WDR4 (Catalog No. ab169526; Abcam), anti-SCLT1 (Catalog No. 14875-1-AP; Proteintech, Rosemont, IL), and Ki67 (Catalog No. GB111499; Servicebio, Wuhan, China) antibodies were prepared following the manufacturer’s instructions.

### Cell lines and reagents

The human lung adenocarcinoma cell lines PC-9 (exon 19 deletion) and HCC4006 (exon 19 deletion) were obtained from the American Type Culture Collection (ATCC). Gefitinib-resistant cells (PC-9/GR and HCC4006/GR) were established by continuously exposing parent cells to increasing gefitinib (Catalog No. ZD1839; Selleck, Shanghai, China) concentrations. Verification of drug-resistant phenotypes was conducted through cell viability assays. PC-9/GR and HCC4006/GR cells were derived from PC-9 and HCC4006 cell lines, respectively, and demonstrated resistance to gefitinib. The cell lines were maintained in RPMI-1640 (Catalog No. 11875093; Gibco, Carlsbad, CA) medium supplemented with 10% fetal bovine serum (FBS) (Catalog No. 16140078; Gibco) and 1% penicillin-streptomycin (PS) (Catalog No. P1400; Solarbio, Beijing, China), and cultured at 37°C in a humidified atmosphere containing 5% CO_2_.

### Cell viability assay

Post various experimental treatments, cell viability was assessed using the CCK-8 Cell Viability Assay Kit (Catalog No. A311-01; Vazyme, Nanjing, China).

### Colony Formation Assay

Cells were cultured until visible cell clones formed, typically taking 7-10 days, with the medium being refreshed every 3 days. Following a rinse with PBS (Catalog No. G4202; Servicebio), the cells are fixed using 4% paraformaldehyde (Catalog No. G1101; Servicebio), and subsequently stained with 1% crystal violet (Catalog No. G1014; Servicebio). The stained cells are rinsed with PBS to remove excess dye, and the plates are air-dried before photographing against a white background.

### Flow cytometry detection

After the cells adhered to the surface of the culture dish, they were transfected with the specified siRNA and subsequently exposed to either DMSO or gefitinib (1 μM) for 72 hours. Apoptosis analyses were conducted following the Annexin V-FITC/PI Apoptosis Detection Kit (Catalog No. A211-02; Vazyme) guidelines.

### Quantitative real-time PCR (qRT-PCR)

Total RNA was extracted using TRIzol reagent (Catalog No. 15596026CN; Invitrogen, Carlsbad, MA), followed by reverse transcription into complementary DNA (cDNA) using the Takara kit (Catalog No. RR047A; Takara, Kusatsu, Japan). Real-time quantitative PCR was conducted with the Takara kit (Catalog No. RR820A; Takara) and QuantStudio 5 Flex real-time qPCR instrument, following the manufacturer’s instructions. GAPDH or β-actin were employed as housekeeping reference genes for normalization. The primer pairs employed for qPCR analysis are detailed below:

METTL1 forward: 5’-GGCAACGTGCTCACTCCAA-3’,
METTL1 reverse: 5’-CACAGCCTATGTCTGCAAACT-3’;
WDR4 forward: 5’-CCACCTCCATAGCAAGCAGTG-3’;
WDR4 reverse: 5’-ACGCTTACTGTCATCGGTTAAAG-3’;
SCLT1 forward: 5’-CATGGAAGTGACTAACCAACAGT-3’;
SCLT1 reverse: 5’-CAGCTCTAATTTGGCTTGCCTA-3’;
GAPDH forward: 5’-GGAGCGAGATCCCTCCAAAAT-3’;
GAPDH reverse: 5’-GGCTGTTGTCATACTTCTCATGG-3’;
β-actin forward: 5’-CATGTACGTTGCTATCCAGGC-3’;
β-actin reverse: 5’-CTCCTTAATGTCACGCACGAT-3’.

### Western blotting analysis

Western blot was conducted as previously described [34]. Primary antibodies used in this investigation are detailed below: anti-METTL1 (Catalog No. ab271063; Abcam), anti-WDR4 (Catalog No. ab169526; Abcam), anti-SCLT1 (Catalog No. 14875-1-AP; Proteintech), anti-Caspase 3 (Catalog No. 19677-1-AP; Proteintech), anti-Cleaved Caspase 3 (Catalog No. 9661; Cell Signaling Technology, Danvers, MA), anti-NF-κB (Catalog No. 8242; Cell Signaling Technology), anti-Phospho-NF-κB (Catalog No. 3033; Cell Signaling Technology), anti-β-actin (Catalog No. GB15001-100; Servicebio) and anti-GAPDH (Catalog No. GB15002-100; Servicebio).

### Dot blot

RNA was purified utilizing the VAHTS mRNA Capture Beads Kit (Catalog No. N401-02; Vazyme). The mRNA underwent decapping with the RNA 5’-pyrophosphohydrolase enzyme (RppH) (Catalog No. M0356S; New England Biolabs, Ipswich, MA). Following denaturation at 65°C for 5 minutes, 500 ng of mRNA was applied onto a Hybond N+ membrane (Catalog No. YA1760; Solarbio). Subsequently, the membrane was UV cross-linked for 5 minutes, blocked with 5% skim milk, and incubated with the m^7^G antibody (Catalog No. ab300740; Abcam). The subsequent step involved incubation with secondary antibodies and the exposure was conducted using a chemiluminescence imaging system. Finally, the bands were immersed in a methylene blue solution (Catalog No. G1300; Solarbio) to visualize the internal reference.

### RNA Interference

Cells were cultured until reaching a density of 30-60%. The transfection was carried out using Lipofectamine RNAiMAX (Catalog No. 13778150; Invitrogen) as per the manufacturer’s guidelines, utilizing either specific siRNA oligonucleotides or through gefitinib treatment. After transfection, cells were incubated for 72 hours and then subjected to qRT-PCR or Western blotting analysis. The siRNA oligonucleotides targeted the following sequences:

siMETTL1: 5’-CCAGCCAUCUUCCGAAGAATT-3’;
siWDR4: 5’-CCUCCUACCUGAAGAAGAATT-3’;
siSCLT1: 5’-UAGGAGAACUAAAUGGGCAGCUGAA-3’.

### Plasmids and transfection

The plasmids shMETTL1, shWDR4, shSCLT1, 3FLAG-METTL1 (NM_005371.6), 3FLAG-METTL1-Mut (E107A, I108A, R109A) and 3FLAG-WDR4 (NM_005228.5) were purchased from GENECHEM (Shanghai, China). Plasmids were transfected into HEK293T cells at 60% confluence using Lipofectamine 3000 (Catalog No. L3000015; Invitrogen) following the manufacturer’s protocol. After a total of 72 hours of incubation, the supernatant was harvested and concentrated using the Lentivirus Concentration Reagent kit (Catalog No. BW-V2001; BIOMIGA, Shanghai, China). Afterward, host cells were infected with the concentrated virus along with 6 μg/mL of polybrene (Catalog No.H8761, Solarbio). Following that, stable polyclonal populations of the transfected cells were chosen by applying either puromycin (Catalog No. P8230, Solarbio) or hygromycin (Catalog No.H8080, Solarbio) over a two-week period, then validated using qRT-PCR or Western blot analysis. The specific shRNA sequences are listed below:

shMETTL1: 5’-CCAGCCATCTTCCGAAGAA-3’;
shWDR4: 5’-CCTCCTACCTGAAGAAGAA-3’;
shSCLT1: 5’-TAGGAGAACTAAATGGGCAGCTGAA-3’.

### RNA stability assay

Cells were subjected to treatment with Actinomycin D (Catalog No. HY-17559; MedChemExpress, Princeton, NJ) for different time intervals: 0, 2, 4, 6, 8, and 10 hours. The levels of target RNA were assessed via qRT-PCR, utilizing GAPDH as the internal control, to determine the relative mRNA abundance at each time point compared to t = 0 hours.

### Animal studies

This investigation utilized female BALB/C nude mice aged five weeks to generate subcutaneous xenograft models. A cell suspension (100 μL, 5×10⁶ cells) was injected into the right axillary region. Tumor growth parameters, specifically longitudinal (a) and transverse (b) diameters, were monitored triweekly, with volumetric calculations performed using the equation V = 1/2·ab². Upon reaching predetermined tumor dimensions, subjects were humanely sacrificed through cervical dislocation, followed by meticulous xenografts excision and cryopreservation in liquid nitrogen.

### Methylated RNA immunoprecipitation (MeRIP)

The isolated mRNA underwent decapping using the RNA 5’-pyrophosphohydrolase enzyme (RppH) (Catalog No. M0356S; New England Biolabs). Subsequently, the mRNA was purified using RNA cleanbeads and cut into fragments of roughly 100 nucleotides in length using RNA Fragmentation Reagents (Catalog No. AM8740, Invitrogen). One-tenth of the decapped mRNA was reserved as input. Following initial processing, residual mRNA was subjected to a 6-hour incubation with anti-m^7^G antibody (Catalog No. RN017M; MBL, Tokyo, Japan) at ambient temperature. The resulting beads-antibody-mRNA conjugates were subsequently subjected to three sequential washes using ice-cold buffer solution. After washing, the immunoprecipitated sample underwent treated with proteinase K, after which RNA was isolated with the aid of glycogen (Catalog No. AM9516; Thermo Fisher Scientific, Grand Island, NY). Finally, quantitative RT-PCR analysis was conducted afterward.

### RNA-seq and RNA-MeRIP-seq

Briefly, total RNA underwent a series of processes including mRNA enrichment, decapping, fragmentation, and purification. Subsequently, cell samples containing 20 ng of mRNA were used to construct RNA-seq libraries using the KAPA Stranded mRNA-seq Kit (Catalog No. KK8421; KAPA, Wilmington, MA). For RNA-MeRIP-seq libraries, the purified decapped mRNA underwent incubation with the anti-m^7^G antibody and was then utilized for library construction. Library sequencing was conducted using paired-end mode on the Illumina HiSeq-PE150 instrument.

### RNA-seq and RNA-MeRIP-seq bioinformatics analyses

RNA-seq bioinformatics analysis was performed as previously described [34]. For RNA-MeRIP-seq bioinformatics analysis, the following steps were conducted: FastQC was used to filter and process RNA-MeRIP-seq reads for each sample. For this, Cutadapt and Trimmomatic were employed to trim 3’ adaptors, eliminate low-quality reads, and obtain high-quality clean reads. The MeRIP library sequences, after being filtered for high-quality reads, were mapped to the UCSC human genome (hg38) using the Hisat2 tool. To pinpoint m^7^G methylation sites within the identified peaks, the MACS2 algorithm was employed for precise detection, and their differences were determined by means of DiffReps.m^7^G motif was detected by the MEME motif analysis with MeRIP-seq data.

### Statistical analysis

Data are presented as mean ± SD unless otherwise specified. Intergroup comparisons were performed using two-tailed Student’s t-tests where applicable. Survival outcomes were assessed through Kaplan-Meier methodology, with statistical significance determined by two-sided log-rank testing. Thresholds for statistical significance were defined as follows: *, *P* < 0.05, **, *P* < 0.01, ***, *P* < 0.001 and ****, *P* < 0.0001.

## Data availability

The raw sequence data from RNA-seq and MeRIP-seq have been deposited in the Genome Sequence Archive [44] in National Genomics Data Center [45], China National Center for Bioinformation / Beijing Institute of Genomics, Chinese Academy of Sciences (GSA-Human: HRA008425). The data is currently confidential and has not yet been released.

## Ethical statement

The present study received approval from the Institutional Review Board of the First Affiliated Hospital of Zhengzhou University (approval number: 2019-KY-0024-001) and adhered to the Helsinki Declaration’s guidelines. All participants gave their explicit written consent, in accordance with institutional policies. The Ethics Committee of the Zhengzhou University granted ethical approval for the animal studies (approval number: ZZU-LAC20220225[01]).

## CRediT author statement

**Shaoxuan Zhou:** Methodology, Validation, Visualization, Writing – original draft. **Yueqin Wang:** Conceptualization, Supervision, Writing – review & editing. **Jingyao Wei:** Formal analysis, Data curation. **Ke An:** Formal analysis, Data curation. **Yong Shi:** Formal analysis. **Xuran Zhang:** Validation, Investigation. **Han Wang:** Validation, Investigation. **Luyao Feng:** Validation, Investigation. **Lulu He:** Validation, Investigation. **Yu Zhang:** Validation, Investigation. **Tong Ren:** Validation, Investigation. **Ouwen Li:** Validation, Investigation. **Yun-Gui Yang:** Conceptualization, Supervision, Project administration, Writing – review & editing. **Quancheng Kan:** Conceptualization, Funding acquisition, Project administration. **Xin Tian:** Conceptualization, Supervision, Funding acquisition, Project administration, Writing – review & editing.

## Acknowledgments

This study was funded by the National Natural Science Foundation of China (32170594), the Zhongyuan Leading Talent in Science and Technology Innovation Program, the Funding for Scientific Research and Innovation Team of The First Affiliated Hospital of Zhengzhou University (ZYCXTD2023010), and the Henan Province University Key Research Project (23A310014).

## Supplementary material

**Figure S1.**
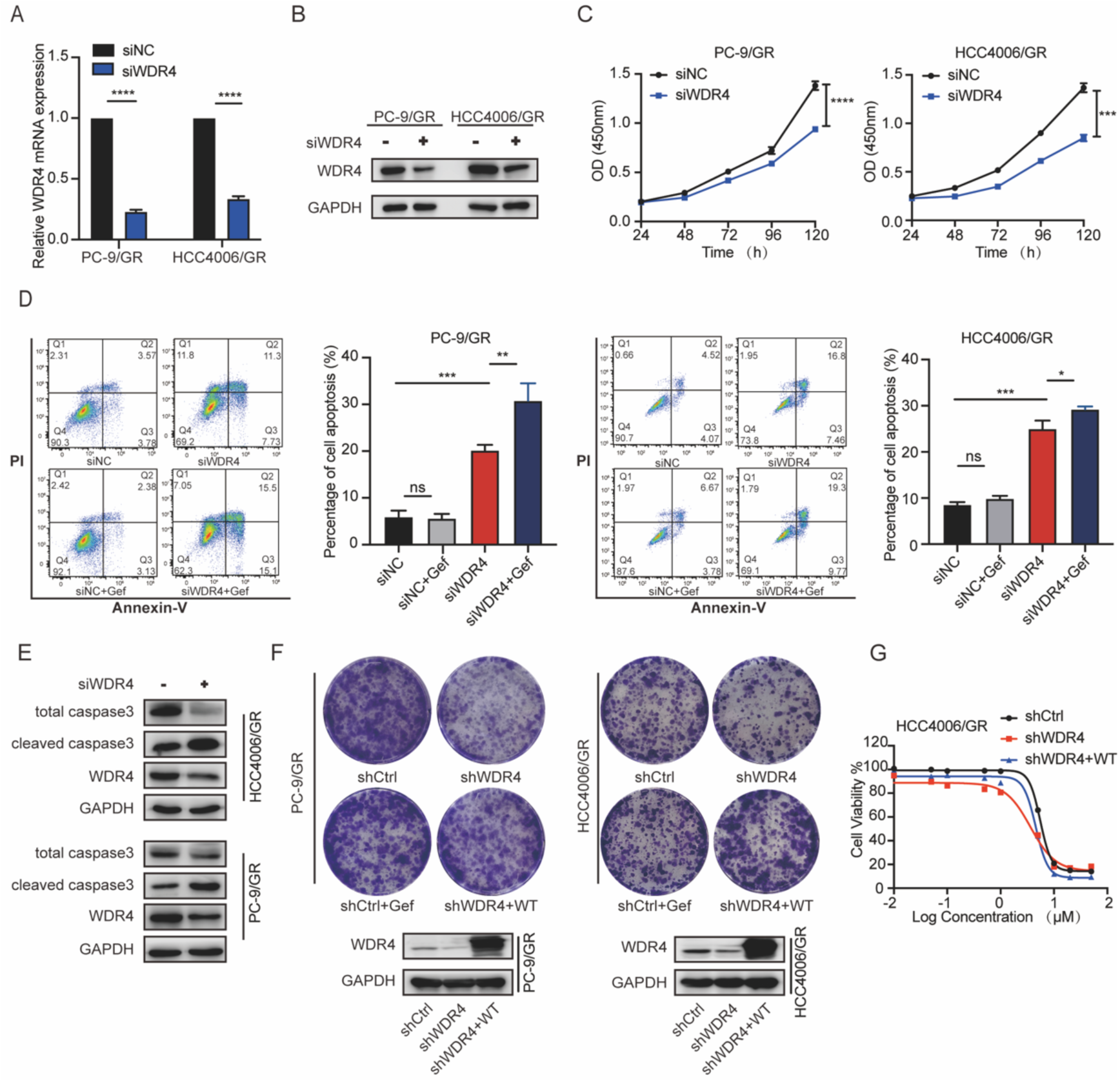
Inhibition of WDR4 overcomes gefitinib resistance *in vitro*. **A.-B.** The mRNA (**A**) and protein expression (**B**) levels of WDR4 in PC-9/GR and HCC4006/GR cells, treated with siRNA for 72 hours, were evaluated using qRT-PCR and Western blotting, respectively. **C.** Knockdown of WDR4 using siRNA was performed in PC-9/GR and HCC4006/GR cells, and the cell viability was assessed using the CCK-8 assay. **D.** PC-9/GR and HCC4006/GR cells transfected with either siNC or siWDR4 were exposed to gefitinib (1 µM) for 72 hours, and cell apoptosis was assessed using flow cytometry. Bar graphs were generated to quantify the data obtained from Annexin V-FITC/PI staining. **E.** The apoptosis marker Cleaved Caspase 3 was detected by Western blotting in PC-9/GR and HCC4006/GR cells treated with siNC or siWDR4 for 72 hours. **F.** The protein expression of WDR4 was assessed in PC-9/GR and HCC4006/GR cells transfected with shWDR4 or WT WDR4 using Western blotting analysis. Additionally, the colony formation ability was evaluated in PC-9/GR and HCC4006/GR using the colony formation assay. **G.** HCC4006/GR cells transfected with shWDR4 or rescue WT WDR4 were treated with varying concentrations of gefitinib for 72 hours. The cell viability was assessed using the CCK-8 assay. Data are shown as mean ± SD. ns, not significant; *, *P* < 0.05; **, *P* < 0.01; ***, *P* < 0.001; ****, *P* < 0.0001. An unpaired t-test was used unless otherwise stated.

**Figure S2.**
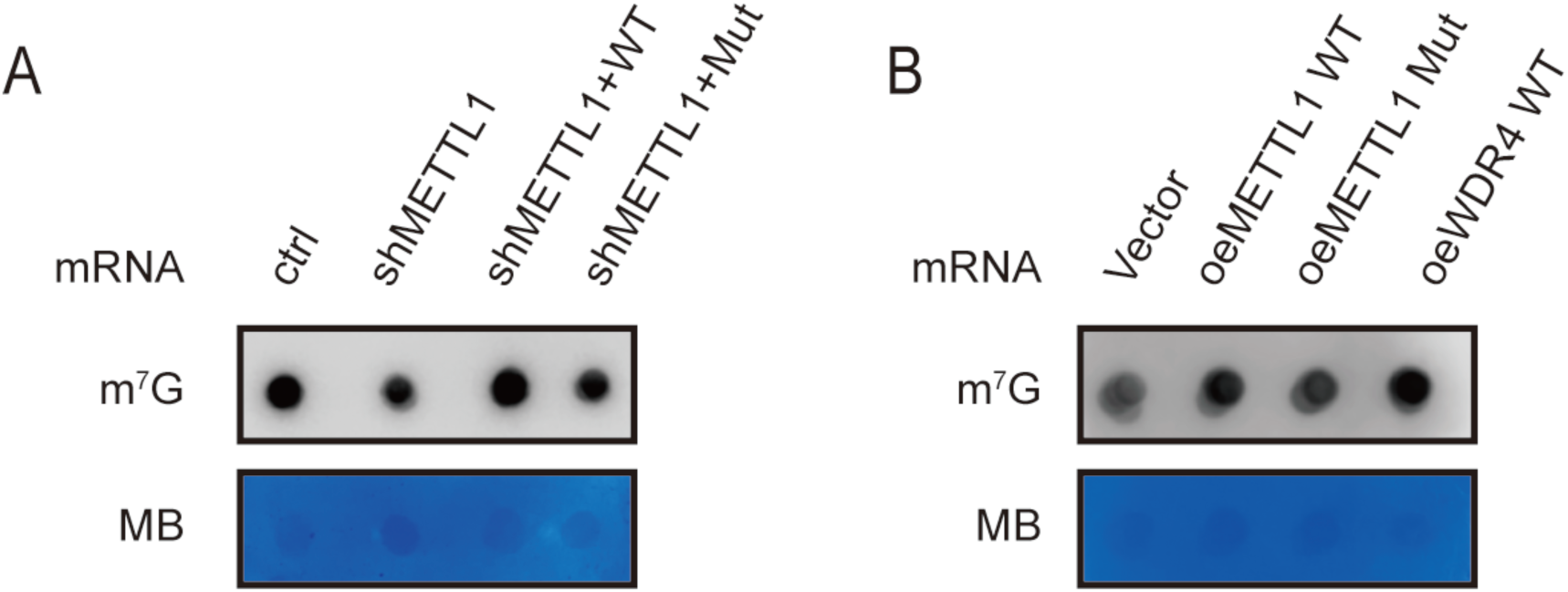
METTL1/WDR4 regulates cellular m^7^G levels in mRNA. **A.** PC-9/GR cells pre-treated with shMETTL1 were stably transfected with either METTL1-WT or METTL1-Mut, and the mRNA m^7^G level was assessed by dot blot assay. **B.** PC-9 cells were stably transfected with either wild-type METTL1/WDR4 or METTL1-Mut, and the mRNA m^7^G level was analyzed by dot blot assay.

**Figure S3.**
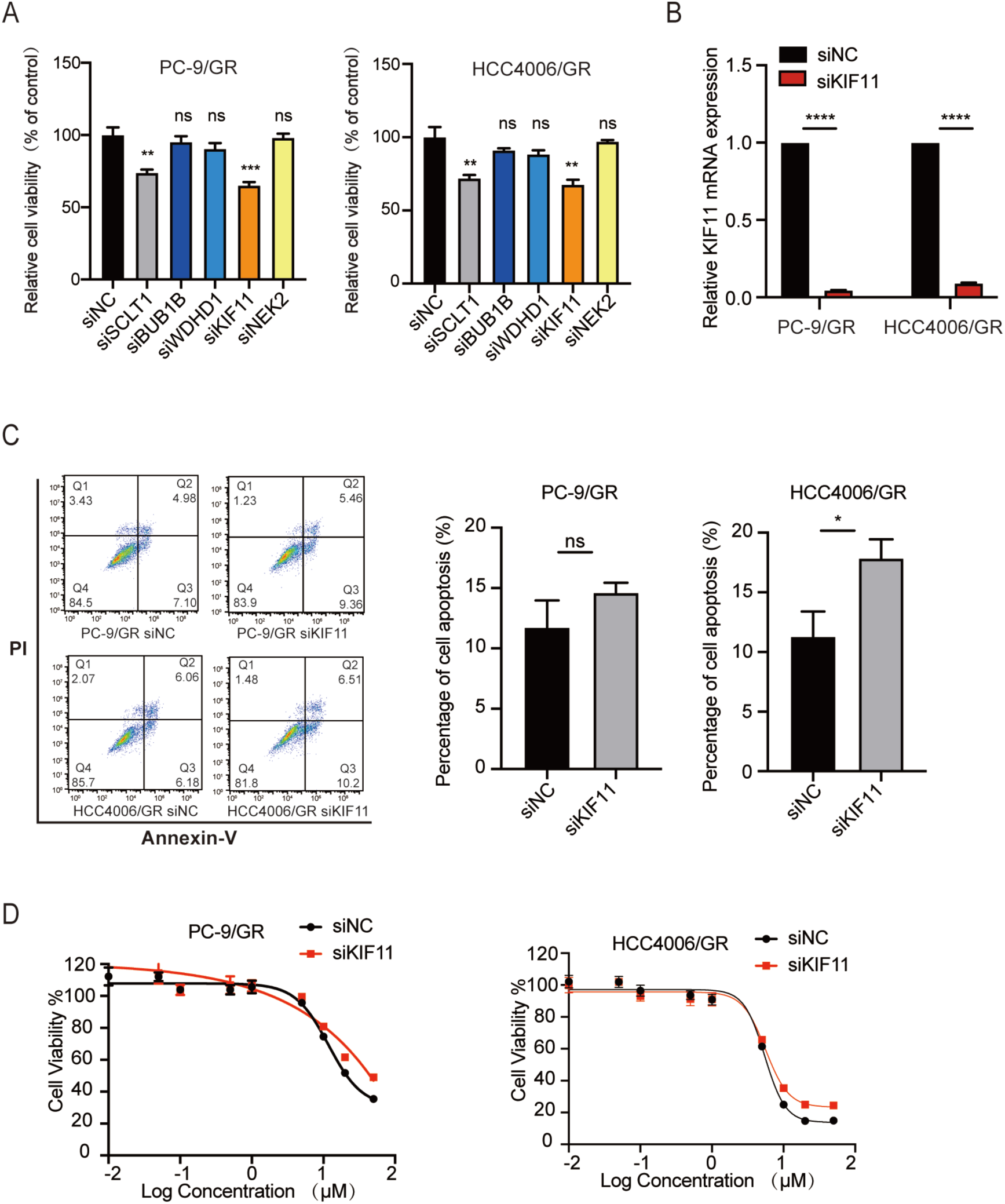
Validation of cell phenotypes following knockdown of genes of interest in gefitinib-resistant cells. **A.** Cell proliferation was evaluated via the CCK-8 assays in PC-9/GR and HCC4006/GR cells transfected with siNC or siRNA for target genes (SCLT1, BUB1B, WDHD1, KIF11, NEK2). **B.** The mRNA levels of KIF11 in PC-9/GR and HCC4006/GR cells, treated with siRNA for 72 hours, were evaluated using qRT-PCR assay. **C.** PC-9/GR and HCC4006/GR cells transfected with either siNC or siKIF11 and cell apoptosis was assessed using flow cytometry. Bar graphs were generated to quantify the data obtained from Annexin V-FITC/PI staining. **D.** PC-9/GR and HCC4006/GR cells transfected with either siNC or siKIF11were treated with varying concentrations of gefitinib for 72 hours. The cell viability was assessed using the CCK-8 assay. Data are shown as mean ± SD. ns, not significant; *, *P* < 0.05; **, *P* < 0.01; ***, *P* < 0.001; ****, *P* < 0.0001. An unpaired t-test was used unless otherwise stated.

**Figure S4.**
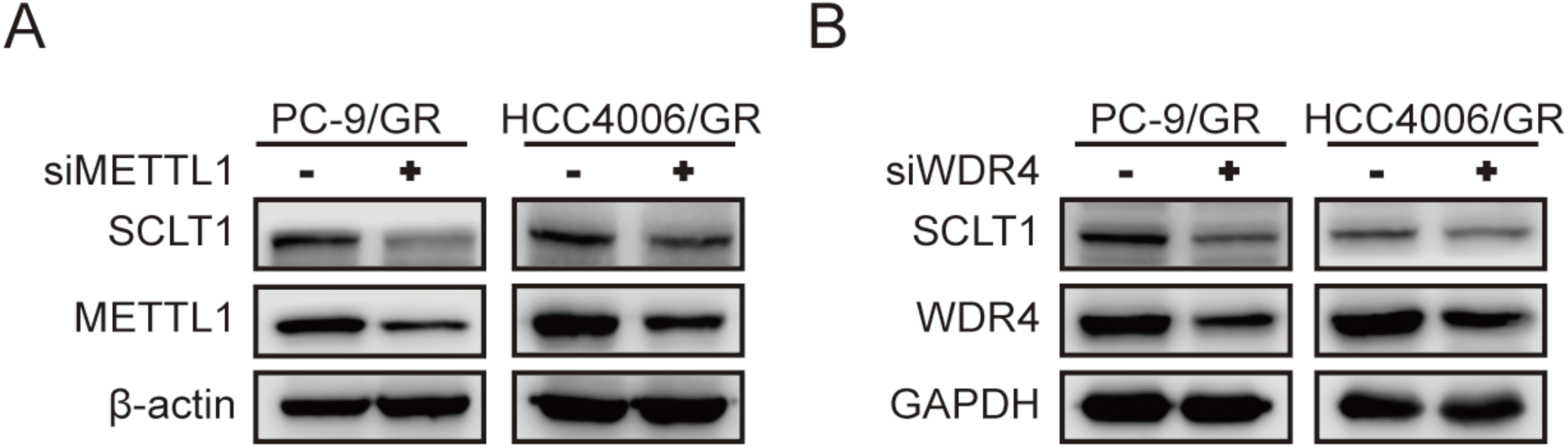
Protein expression levels of SCLT1 following METTL1 and WDR4 knockdown. **A.-B.** The protein expression levels of SCLT1 were evaluated via Western blotting in PC-9/GR and HCC4006/GR cells treated with siMETTL1 (**A**) or siWDR4 (**B**) for 72 hours.

**Figure S5.**
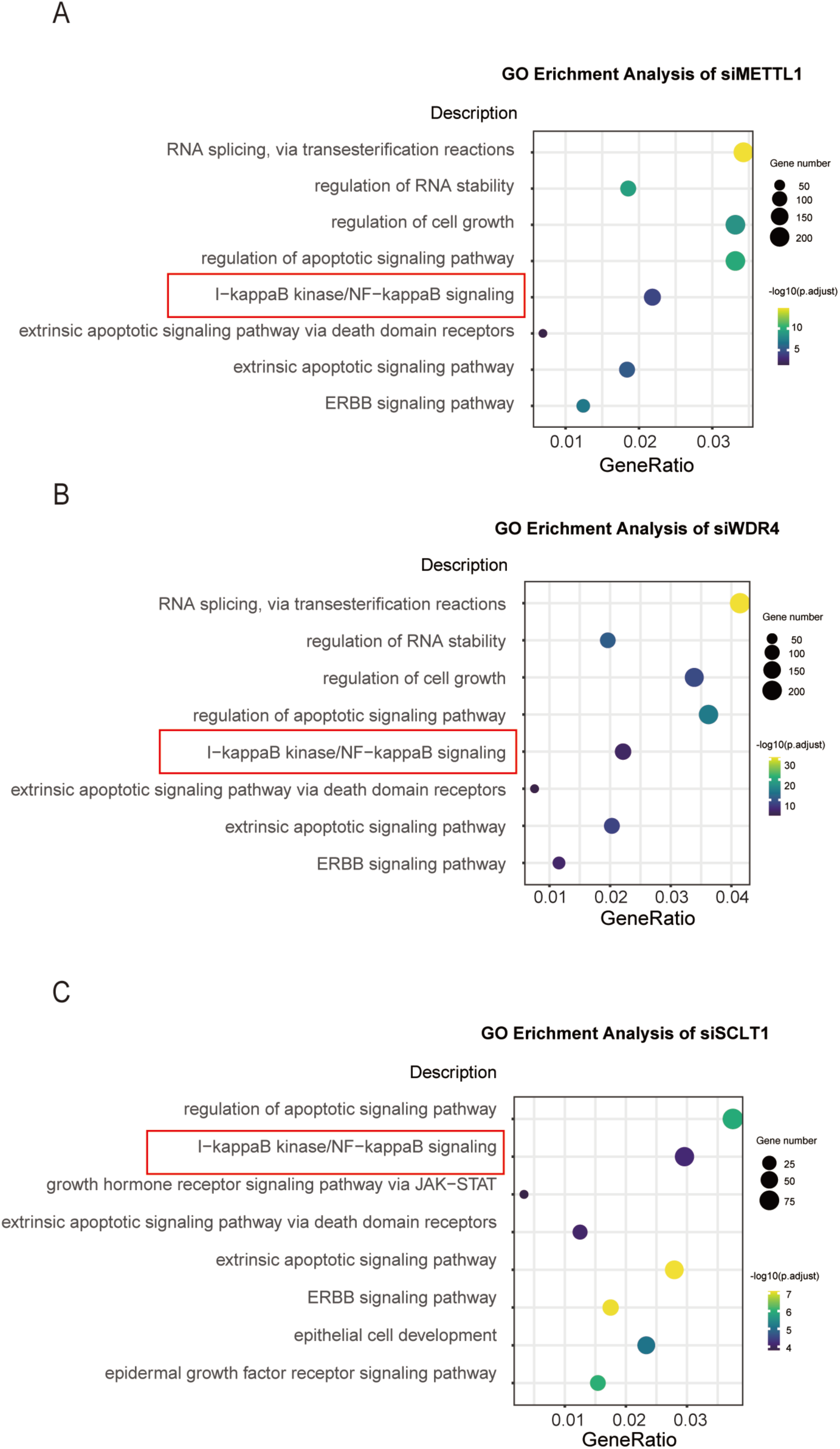
m^7^G modification regulates gefitinib resistance through the METTL1/WDR4-SCLT1-NF-κB axis. **A.-C.** GO pathway analysis of mRNA expression changes in PC-9/GR cells post METTL1 (**A**), WDR4 (**B**) and SCLT1 (**C**) knockdown.

**Table S1 Summary of RNA-seq Gene data**

**Table S2 Summary of m^7^G Methylated RNA sites data**

